# Molecular Surveillance Identifies Evidence of Getah Virus (GETV) in Mosquito Vectors in Alabama, USA

**DOI:** 10.64898/2025.12.04.692395

**Authors:** Ting Li, Tafarah McDaniel, Qiana Matthews, Efa Barnard, Rachana Pandit, Hongzhuan Wu

## Abstract

The Getah virus (GETV) is a mosquito-borne RNA virus in the Togaviridae family and the Alphavirus genus, associated with severe disease outbreaks in livestock across Asia-Pacific regions. Since its first isolation in Malaysia in 1955, GETV has been reported in Eurasia and the South Pacific, yet documented cases in the United States remain exceptionally rare, leaving major gaps in regional vector competence and surveillance data. To evaluate the potential of local mosquito populations as GETV carriers, field collections were conducted in Montgomery and Pike Road neighborhoods of Alabama from 2023 to 2024, capturing multiple mosquito species for molecular screening. Initial real-time qPCR assays on 200 pooled and individual samples suggested that approximate 30% of specimens demonstrated possible GETV carriage potential. To validate viral presence, two distinct primer sets were designed to amplify viral genome fragments. Mosquito RNA was reverse-transcribed into cDNA, followed by conventional PCR. The first PCR, targeting ∼280 bp, produced single or multiple amplicon bands in 32% of samples via gel electrophoresis. These positive cDNAs were re-amplified with a second primer pair targeting a ∼430 bp fragment from a separate genomic region, yielding confirmatory bands in 30% of specimens. Amplified products were purified and Sanger sequenced, revealing approximate 95% nucleotide similarity to the wild-type GETV reference genome. This study delivers the first field-based molecular evidence supporting GETV detection in Alabama mosquitoes, signaling either localized emergence or potential introduction of the virus. These findings underscore the need for expanded vector competence profiling, arboviral surveillance, and livestock disease preparedness in the southeastern United States, strengthening both public health readiness and state-level vector monitoring strategies.

## 1. Introduction

Mosquitoes are often regarded as the leading transmitters of diseases to humans due to their role in spreading various pathogens. For example, malaria, transmitted by Anopheles mosquitoes, results in over half million deaths annually with most cases occurring in Africa [1–2]. West Nile Virus spread by Culex mosquitoes [3–4] and Yellow Fever transmitted by Aedes mosquitoes [5] have been reported in the United States and Worldwide. Monitoring mosquito populations is a crucial tool for understanding their distribution and plays a pivotal role in preventing the transmission of mosquito-borne diseases [6–7]. Getah virus (GETV) is an enveloped, single-stranded, positive-sense RNA virus with a genome of approximately 11,000–12,000 nucleotides [8]. The virus was first isolated in 1955 from a Culex mosquito in Malaysia, and has since been detected in multiple mosquito species, livestock, and wild animals, demonstrating a broad host range and diverse tissue tropisms [8]. Although no confirmed human clinical illness attributed to GETV has been reported to date [9], the virus is a well-recognized veterinary pathogen with substantial impact on livestock health. GETV infection in pigs can progress rapidly, with reports showing that in one outbreak, ∼200 piglets died within 5–10 days postpartum [10]. Viral RNA has also been identified in the brains of deceased piglets and in multiple porcine organs, supporting both potential neuroinvasion and multi-organ replication, as well as transplacental infection consistent with vertical transmission. GETV exhibits wide tissue dissemination, indicating the ability to infect reproductive, neural, and systemic organ systems [10]. Infected horses may develop fever, systemic rash, and limb edema, while pigs, particularly neonates and pregnant sows, can experience piglet mortality, abortion, fetal mummification, stillbirth, and reproductive disorders, contributing to considerable agricultural and economic loss [11]. GETV is maintained through a mosquito–vertebrate transmission cycle, in which horses and domestic pigs serve as major viral amplifiers [12]. Vertical transmission is suspected in pigs, likely occurring via placental transfer [13], whereas in horses, direct contact and mosquito exposure are regarded as primary transmission routes [14]. Geographically, GETV is distributed across a vast region spanning Malaysia, mainland China, Russia, and parts of southern Russia, where it infects horses, pigs, cattle, poultry, wildlife, and multiple mosquito vectors [8]. Its ecological transmission pattern resembles that of Japanese encephalitis virus, another Culex-transmitted arbovirus amplified in pigs, underscoring shared epidemiological risk landscapes, vector overlap, and livestock vulnerability in Asia [14]. Over the past decade, frequent epidemics in countries such as China, Japan, India, and South Korea have elevated GETV as a regional priority for vector-borne livestock disease surveillance and control [15]. Collectively, current evidence supports GETV as a high-risk, widely spreading arboviral pathogen of veterinary relevance, with increasing concern for its transmission potential, evolving viral variants, reproductive and neural pathogenicity in mammals, and geographic expansion across Asia-Pacific regions.

Interestingly, previous studies found that GETV also holds potential as an oncolytic agent in cancer research. Oncolytic viruses are recognized for their ability to specifically target and destroy tumor cells while sparing healthy tissues. Several viruses including herpes simplex virus, vaccinia virus, retrovirus, vesicular stomatitis virus, and adenovirus have been developed or engineered as oncolytic agents [16]. These viruses are designed to exploit the genetic abnormalities of cancer cells, which leads to their selective replication and destruction. Oncolytic viruses initiate an immune response by triggering local inflammation within the tumor microenvironment. For instance, Alphavirus M1 demonstrates potent antitumor activity via intravenous infusion although its replication can be limited by the zinc-finger antiviral protein (ZAP). Oncolytic viruses can either occur naturally or be genetically modified to replicate specifically within cancer cells, leveraging tumor-specific promoters or deletions in crucial viral genome regions [17]. This strategy has garnered significant attention over the years. Advances in scientific research have led to substantial improvements in the design and application of these viral agents. Through genetic engineering, researchers can now tailor oncolytic viruses to enhance the specificity for cancer cells and boost their replicative capacity.

Studies have shown that GETV can be amplified in mosquitoes through a simple bite with infected animals acting as reservoirs from which uninfected mosquitoes can acquire the virus [18]. GETV is related to other alphaviruses such as Chikungunya virus (CHIKV) and Ross River virus (RRV), which are pathogenic to humans. Their resistance partly drives the spread of these mosquitoes to new regions where global travel is increased, which complicates effective control measures and facilitates their spread to new areas [19]. Regarding the distribution of GETV, 129 strains of the virus were listed in GenBank with an additional 41 strains identified in Chinese and English references as of March 2022. Of these, only 24 strains had been isolated. In respect to host animals, 56 strains of GETV had been isolated from various species by March 2022, which included pigs (37 strains), horses (12 strains), wild boars (2 strains), red pandas (2 strains), cattle (1 strain), blue fox (1 strain) and a fox (1 strain) [8]. The distribution of GETV has been extensively studied in China, Mongolia, Japan, South Korea, India, Thailand, and other regions across Asia [8]. Continuous monitoring of the virus in these animals as well as in mosquitoes is essential for detecting potential illnesses and preventing transmission to avert outbreaks. Our study aims to assess the potential prevalence of Getah virus (GETV) transmission among mosquito populations in Alabama. We conducted field-based mosquito surveillance and laboratory identification of species, confirming the presence of *Culex quinquefasciatus*, *Aedes albopictus*, *A. triseriatus*, and *A. japonicus*. RNA was extracted from 200 mosquito pools and individual adult specimens and analyzed for GETV infection using real-time qPCR, followed by viral genome PCR amplification and sequencing for confirmation. Preliminary results indicate that approximately 30% of the tested samples yielded positive amplification signals consistent with GETV, suggesting possible circulation of GETV-like viral RNA within local mosquito populations.

## Materials and Methods

### 1. Mosquito collection and identification

Mosquito samples were collected from multiple locations in the Montgomery and Pike Road neighborhoods of Alabama during 2023–2024 as part of ongoing mosquito population surveillance. Adult mosquitoes were captured using CDC gravid traps (Model 1712) and CDC light traps [20–21]. The gravid trap is specifically designed to target Culex species [22]; it uses a pan containing hay infusion and dry ice to attract gravid females seeking oviposition sites, after which an upward airflow generated by an internal fan directs the mosquitoes into a collection bag. CDC light traps, operated with or without the light source, attracted mosquitoes through a combination of illumination and airflow. All traps were assembled in the laboratory and deployed weekly in selected urban residential areas in Montgomery and Pike Road, AL. Mosquitoes were retrieved the following day and transported to the laboratory for identification. Species identification was performed using standard morphological keys from “Identification and Geographical Distribution of the Mosquitoes of North America, North of Mexico” [23]. The total number of collected mosquitoes and species composition were recorded. Fully developed adult mosquitoes were stored at –80°C to preserve RNA integrity for subsequent molecular analyses. Male mosquitoes were discarded, and only female mosquitoes were retained for downstream testing.

### 2. RNA isolation and cDNA preparation

Total RNA was extracted from individual female mosquito samples using TRIzol Reagent (Invitrogen) following the manufacturer’s standard protocol [24]. RNA concentration and purity were assessed using a NanoDrop spectrophotometer prior to cDNA synthesis. cDNA was generated from equal amounts of RNA using the Verso cDNA Synthesis Kit (Thermo Scientific) according to the manufacturer’s instructions, and the resulting cDNA concentrations were measured and recorded using a NanoDrop spectrophotometer. All cDNA samples were stored at −80°C until use in real-time qPCR and conventional PCR amplification. For positive controls, the standard Getah virus (GETV) strain (ATCC VR-369) was obtained from the American Type Culture Collection. Viral RNA was isolated using the QIAamp Viral RNA Mini Kit (Qiagen) following the manufacturer’s protocol. RNA concentration and purity were evaluated prior to downstream cDNA synthesis to ensure suitability for assay optimization and validation.

### 3. Quantitative real-time PCR (qRT-PCR)

To detect Getah virus (GETV) in mosquito samples, specific qRT-PCR primers were designed and synthesized by Integrated DNA Technologies (IDT) based on the GenBank reference genome sequence of GETV (NC_006558.1). The primer sequences used were: GETV-F, 5′-TCGCAAGCTACACCACATAG-3′ and GETV-R, 5′-CATACTCTGTCTCTGCCCTTTC-3′. qRT-PCR was performed using the ABsolute QPCR SYBR Green ROX Mix (Thermo Fisher Scientific) on a CFX96 Real-Time System (C1000 Thermal Cycler, Bio-Rad). Each 15-µl reaction contained 1× SYBR Green master mix, 1 µl of cDNA template, and GETV-specific primers at a final concentration of 3–5 µM. cDNA synthesized from purified GETV RNA served as the positive control, whereas no-template controls (NTCs) were included as negative controls. All reactions were performed in triplicate. The thermal cycling conditions consisted of an initial enzyme activation at 95°C for 15 min, followed by 40 cycles of 95°C for 15 s, 60°C for 30 s, and 72°C for 30 s. Ct values were recorded for each reaction well.

To determine the optimal Ct threshold for distinguishing true positive amplification from background signal, a receiver operating characteristic (ROC) curve analysis was conducted. Serial dilutions of purified GETV RNA (10³, 10², 20, and 1 copy per reaction) were used as true positives, and two NTC reactions served as true negatives. Ct values were used as continuous predictors for assay performance. ROC analysis was performed using Python’s scikit-learn library, with sensitivity (true positive rate) and 1 – specificity (false positive rate) calculated across all potential Ct cutoffs. The optimal diagnostic threshold was defined using Youden’s Index (J = sensitivity + specificity – 1), and the area under the ROC curve (AUC) was calculated to evaluate overall assay accuracy [25].

### 4. Polymerase Chain Reaction and sequencing analysis

To further confirm the presence of GETV in mosquito samples, conventional PCR was performed to amplify two regions of the viral genome, generating ∼280 bp and ∼430 bp products from mosquito cDNA and from the positive control. Reactions were carried out using Phusion High-Fidelity PCR Master Mix (Thermo Scientific) following the manufacturer’s protocol. The thermal cycling conditions consisted of an initial denaturation at 98°C for 30 s, followed by 40 cycles of 98°C for 15 s, 60°C for 30 s, and 72°C for 30 s, with a final extension step at 72°C for 7 min. Primers were designed based on the GETV reference genome (NC_006558.1) as follows: GETVF1, 5′-CGACCGAGCGATAGAGTAATG-3′ and GETVR1, 5′-TAGGTACAGACCGGGAAAGA-3′; GETVF2, 5′-CAGGTGGATGTAGAGGAACTAAC-3′ and GETVR2, 5′-CATACTCTGTCTCTGCCCTTTC-3′. Agarose gel electrophoresis (1% agarose) was used to visualize PCR products, with a GeneRuler 100 bp DNA ladder (Thermo Scientific) serving as the size marker. Single, distinct amplification bands matching the expected sizes of the positive control were excised and purified using the PureLink Quick Gel Extraction and PCR Purification Combo Kit (Invitrogen). Purified products were submitted to Eurofins Genomics LLC for Sanger sequencing. Sequence data were analyzed using NCBI BLASTn to confirm GETV identity and determine sequence similarity.

## Results

### 1. Mosquito samples, RNA and cDNA quality and quantity

Mosquitoes were collected from multiple neighborhoods in Montgomery and Pike Road, Alabama, during the summer and fall of 2023 and 2024. The species identified included *Aedes albopictus*, *A. triseriatus*, *A. japonicus*, and *Culex quinquefasciatus*. A total of 200 adult mosquito samples—comprising both pooled samples and individual mosquitoes—were processed for the detection of GETV. Of these, 130 samples originated from Pike Road and 70 from Montgomery (Table 1). Total RNA were extracted from each sample, and RNA was subsequently reverse transcribed into cDNA for downstream real-time PCR analysis and conventional PCR reactions. RNA and cDNA concentrations, along with A260/280 ratios, were measured to assess sample purity. RNA yields were generally high in mosquito pool samples and showed expected variation among individual mosquitoes, whereas cDNA concentrations were relatively consistent across samples. Both RNA and cDNA demonstrated high purity and quality, supporting their suitability for molecular detection assays.

**Table 1.**
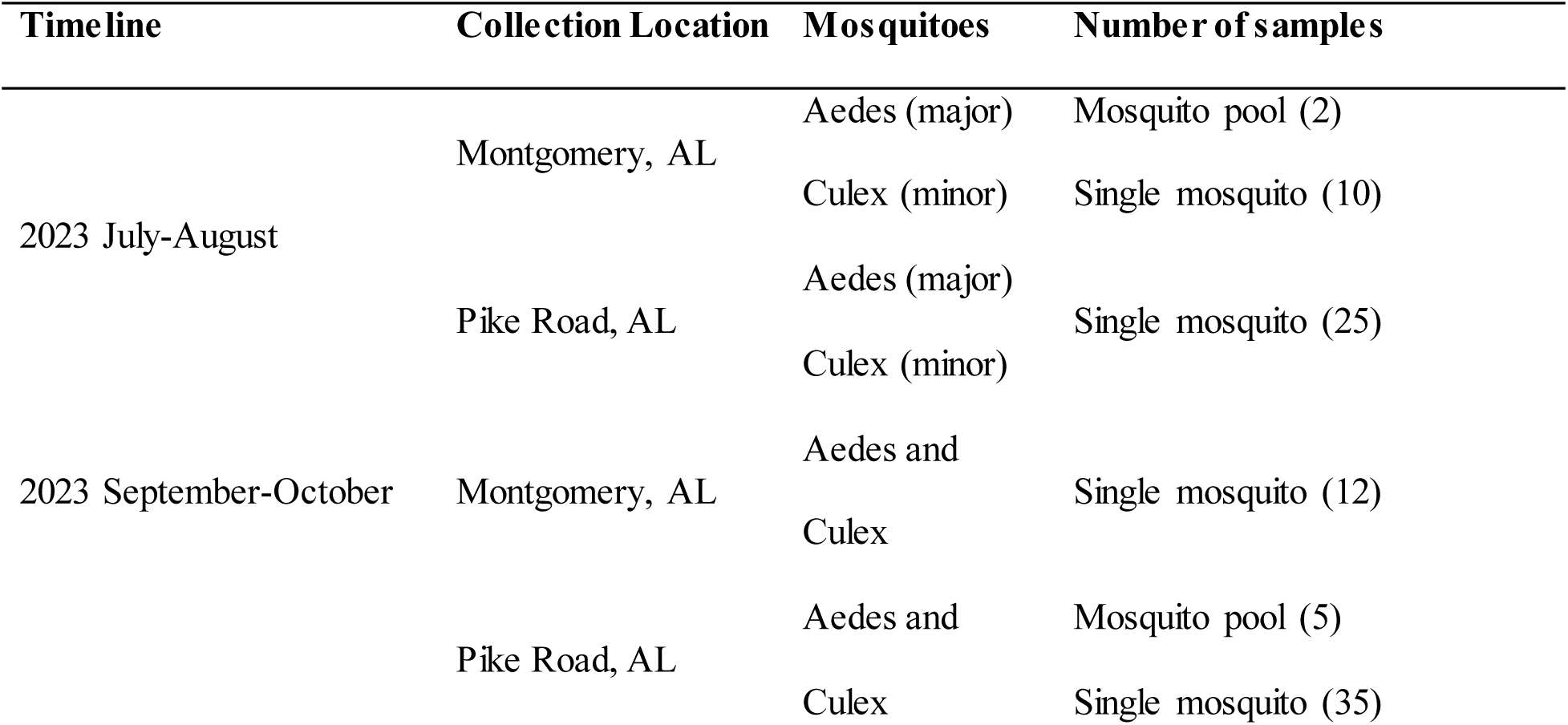
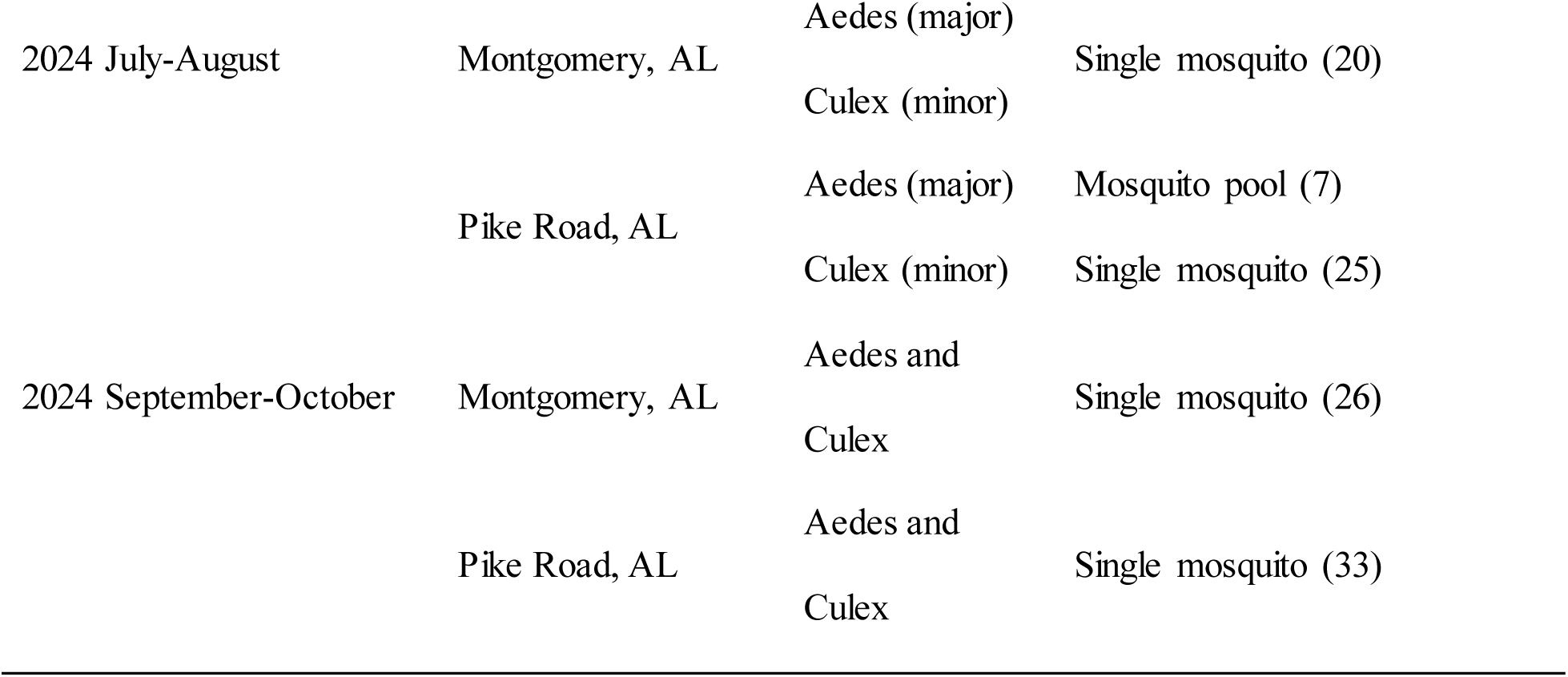
Mosquito Collection and Species Identification.

| Timeline | Collection Location | Mosquitoes | Number of samples |
| --- | --- | --- | --- |
| 2023 July-August | Montgomery, AL | Aedes (major) | Mosquito pool (2) |
|  |  | Culex (minor) | Single mosquito (10) |
|  | Pike Road, AL | Aedes (major) | Single mosquito (25) |
|  |  | Culex (minor) |  |
| 2023 September-October | Montgomery, AL | Aedes and | Single mosquito (12) |
|  |  | Culex |  |
|  | Pike Road, AL | Aedes and | Mosquito pool (5) |
|  |  | Culex | Single mosquito (35) |
| 2024 July-August | Montgomery, AL | Aedes (major) | Single mosquito (20) |
|  |  | Culex (minor) |  |
|  | Pike Road, AL | Aedes (major) | Mosquito pool (7) |
|  |  | Culex (minor) | Single mosquito (25) |
| 2024 September-October | Montgomery, AL | Aedes and<br>Culex | Single mosquito (26) |
|  | Pike Road, AL | Aedes and<br>Culex | Single mosquito (33) |

### 2. Quantitative Real-time PCR for GETV identification

To evaluate the presence of GETV in mosquito samples, quantitative real-time PCR was performed using SYBR Green chemistry and cDNA templates. Serial dilutions of purified GETV RNA yielded Ct values ranging from 20.07 to 37.84 across decreasing template concentrations, whereas negative controls amplified only at Ct = 40. ROC curve analysis demonstrated excellent discriminatory performance of the qPCR assay, with an AUC of 1.00, indicating perfect separation of positive and negative samples within the tested dataset. Youden’s Index identified an optimal diagnostic cutoff of Ct = 37.84, which maximized combined sensitivity and specificity for viral RNA detection. A **l** reactions with Ct ≤ 37.84 were correctly classified as positive, while all negative controls exceeded this threshold. This cutoff therefore represents the upper limit at which true viral amplification can be reliably distinguished from background noise under our assay conditions, and it is consistent with previously reported thresholds in the literature [26].

PCR amplification using GETV-specific primers was monitored in real-time, and each mosquito sample was analyzed in triplicate. Mean Ct values were calculated for all samples. Based on both the literature and our ROC-derived cutoff, a Ct threshold of 37 was used to determine GETV positivity. Samples with mean Ct values ≤ 37 were classified as positive, whereas samples with no detectable amplification were recorded as “N/A.” Standard GETV RNA used as a positive control showed Ct values between 20 and 25, confirming assay sensitivity. Approximately 30% of mosquito samples exhibited Ct values between 33 and 37, indicating detectable levels of GETV-like RNA in both Montgomery and Pike Road mosquito populations (Table 2).

**Table 2.** Quantitative Real-time PCR for GETV Identification.

| Timeline | Collection Location | Sample type | # of samples | Ct mean |
| --- | --- | --- | --- | --- |
| 2023 July-August | Montgomery, AL | Single females | 2 | 33.51~34.28 |
|  | Pike Road, AL | Single females | 10 | 35.12 ~ 36.58 |
| 2023 September-October | Montgomery, AL | Single females | 1 | 35.63 |
|  | Pike Road, AL | Pool and single females | 15 | 35.62 ~ 36.11 |
| 2023 July-August | Montgomery, AL | Single females | 5 | 33.93 ~ 36.05 |
|  | Pike Road, AL | Pool and single females | 8 | 33.48 ~ 34.64 |
| 2023 September-October | Montgomery, AL | Single females | 8 | 35.24 ~ 36.73 |
|  | Pike Road, AL | Single females | 11 | 34.69 ~ 36.31 |

### 3. PCR and sequencing analysis

Approximately 30% of the RNA samples produced PCR amplification bands of the expected size corresponding to the positive control. These amplified products were subsequently purified and submitted for Sanger sequencing. The resulting sequence chromatograms were processed and analyzed by aligning them to the reference viral sequence used as the positive control. Sequence identity among the amplified products ranged from 95% to 96%, confirming that the detected amplicons were derived from the target viral RNA rather than nonspecific amplification. This level of sequence similarity indicates successful detection of the virus in a subset of field samples, although minor base variations may reflect natural genetic diversity, sequencing artifacts, or low-template amplification effects.

## Conclusion and Discussion

Mosquito samples for this study were collected from various neighborhoods in Montgomery and Pike Road, Alabama. These species are significant vectors of various pathogens. For example, *Culex quinquefasciatus* is a primary vector of West Nile Virus, filarial worms, and avian malaria parasites [27], while *Aedes albopictus* is a key vector for Zika, dengue, chikungunya, and yellow fever [28]. The distribution of Culex and Aedes species extends across the southeastern United States [29–30]. The Getah virus (GETV), first identified in Culex mosquitoes in Malaysia in 1955 [31], has since been isolated from other mosquito species and hosts, primarily in Japan [32–33] and China [34]. The purpose of highlighting GETV is to raise awareness about the global impact of mosquito-borne viruses. This virus has caused outbreaks in swine, leading to mild to severe symptoms in livestock such as horses, cattle, pigs, poultry, and other wild animals [35]. However, no clinical symptoms have been reported in humans from GETV infections [36]. GETV has a linear, positive-sense, single-stranded RNA genome ranging from 11 to 12 kb in size [37–38], mutates rapidly, and is transmitted by various mosquito species. The primary vectors include Culex, Aedes, and Anopheles, depending on the region. Although there have been no reports of GETV in the Americas, the distribution of alphaviruses is expanding due to the increasing mosquito populations, adaptation to new vectors, and growing international travel [39–40]. GETV transmission is seasonal, and the virus may be reintroduced into temperate climates from southern regions each year as mosquito vectors migrate. Monitoring mosquito populations helps identify transmission hotspots and assess the effectiveness of control measures [41]. There is also evidence that human-driven land use changes can directly and indirectly influence the transmission dynamics of vector-borne pathogens [42–44]. In our mosquito sample collection, 70% were identified as Aedes species, while 30% were Culex mosquitoes. Our hypothesized that the mosquitoes collected in Alabama may carry the GETV, potentially due to factors such as climate change, global warming, and urbanization. To test this hypothesis, we employed molecular techniques to determine the prevalence of GETV in the samples. In this study, 200 individual mosquitoes and mosquito pools were tested for GETV infection. The real-time qPCR results indicate that approximately 30% of the mosquitoes tested positive for GETV, based on a cutoff threshold of 37 cycles. Standard GETV RNA was used as a positive control. To further confirm the real-time PCR results, the PCR was run to amplify for two different segments of the viral genome, showing that ∼30% mosquito samples may carry the GETV. In this study, 65% of the total samples were collected from the Pike Road neighborhood, which is located near a farm, while 35% came from three separate locations in Montgomery, including one near Jefferson Hospital downtown. While human infections with GETV are currently rare, the potential for outbreaks should not be underestimated, especially given the changing environmental conditions and the impact of climate change on mosquito populations. Addressing the threat of vector-borne diseases requires robust surveillance systems, effective interventions, and public health strategies, including mosquito population monitoring, infection rate assessments, and preventative measures such as vaccine development and public awareness campaigns. Effective surveillance of mosquito vectors is essential for public health management and control of vector-borne diseases. By employing a variety of methods, from traditional entomological surveys to modern molecular techniques, public health authorities can better understand and respond to mosquito distribution patterns.

## Acknowledgement

This work was supported by a pilot mechanism through an institutional development grant (Award number: DICRIDG-22-1037199-01-DICRIDG) from the American Cancer Society to Alabama State University.

## Reference

1. WHO, Malaria, 2025. https://www.who.int/health-topics/malaria?utm_source=chatgpt.com#tab=tab_1. Access on December, 01, 2025.

2. CDC, Fighting the World’s Deadliest Animal, Aug. 12, 2025. https://www.cdc.gov/global-health/impact/fighting-the-worlds-deadliest-animal.html?utm_source=chatgpt.com. Access on December, 01, 2025.

3. CDC, About West Nile, Sept. 10, 2025. https://www.cdc.gov/west-nile-virus/about/index.html?utm_source=chatgpt.com. Access on December, 01, 2025.

4. Hayes, E. B., Komar, N., Nasci, R. S., Montgomery, S. P., O’Leary, D. R., & Campbell, G. L. (2005). Epidemiology and transmission dynamics of West Nile virus disease. Emerging infectious diseases, 11(8), 1167.

5. CDC, Yellow Fever: Causes and How It Spreads, May 15, 2024. https://www.cdc.gov/yellow-fever/causes-and-spread/index.html?utm_source=chatgpt.com. Access on December, 01, 2025.

6. Wilson, A. L., Courtenay, O., Kelly-Hope, L. A., Scott, T. W., Takken, W., Torr, S. J., & Lindsay, S. W. (2020). The importance of vector control for the control and elimination of vector-borne diseases. PLoS neglected tropical diseases, 14(1), e0007831.

7. Brown, H. E., Sedda, L., Sumner, C., Stefanakos, E., Ruberto, I., & Roach, M. (2021). Understanding mosquito surveillance data for analytic efforts: a case study. Journal of medical entomology, 58(4), 1619–1625.

8. Li, B., Wang, H., & Liang, G. (2022). Getah virus (Alphavirus): an emerging, spreading zoonotic virus. Pathogens, 11(8), 945.

9. Wang, A., Zhou, F., Liu, C., Gao, D., Qi, R., Yin, Y., et al. (2022). Structure of infective Getah virus at 2.8 Å resolution determined by cryo-electron microscopy. Cell Discovery, 8(1), 12.

10. Yang, T., Li, R., Hu, Y., Yang, L., Zhao, D., Du, L., et al. (2018). An outbreak of Getah virus infection among pigs in China, 2017. Transboundary and emerging diseases, 65(3), 632–637.

11. Liu, H., Zhang, X., Li, L. X., Shi, N., Sun, X. T., Liu, Q., et al. (2019). First isolation and characterization of Getah virus from cattle in northeastern China. BMC veterinary research, 15(1), 320.

12. Swine Health Information Center. Getah Virus Fact Sheet. 2021.

13. Wu, Y., Gao, X., Kuang, Z., Lin, L., Zhang, H., Yin, L., et al. (2024). Isolation and pathogenicity of a highly virulent group III porcine Getah virus in China. Frontiers in Cellular and Infection Microbiology, 14, 1494654.

14. Azerigyik, F. A., Faizah, A. N., Kobayashi, D., Amoa-Bosompem, M., Matsumura, R., Kai, I., et al. (2023). Evaluating the mosquito host range of Getah virus and the vector competence of selected medically important mosquitoes in Getah virus transmission. Parasites & Vectors, 16(1), 99.

14. Zhang, Y., Li, Y., Guan, Z., Yang, Y., Zhang, J., Sun, Q., et al. (2022). Rapid differential detection of Japanese encephalitis virus and Getah virus in pigs or mosquitos by a duplex TaqMan real-time RT-PCR assay. Frontiers in Veterinary Science, 9, 839443.

15. Shi, N., Zhu, X., Qiu, X., Cao, X., Jiang, Z., Lu, H., & Jin, N. (2022). Origin, genetic diversity, adaptive evolution and transmission dynamics of Getah virus. Transboundary and emerging diseases, 69(4), e1037–e1050.

16. Lin, D., Shen, Y., & Liang, T. (2023). Oncolytic virotherapy: basic principles, recent advances and future directions. Signal transduction and targeted therapy, 8(1), 156.

17. Lin, Y., Zhang, H., Liang, J., Li, K., Zhu, W., Fu, L., et al. (2014). Identification and characterization of alphavirus M1 as a selective oncolytic virus targeting ZAP-defective human cancers. Proceedings of the National Academy of Sciences, 111(42), E4504–E4512.

18. Sam, S. S., Teoh, B. T., Chee, C. M., Mohamed-Romai-Noor, N. A., Abd-Jamil, J., Loong, S. K., et al. (2018). A quantitative reverse transcription-polymerase chain reaction for detection of Getah virus. Scientific reports, 8(1), 17632.

19. Powers, A. M., Brault, A. C., Shirako, Y., Strauss, E. G., Kang, W., Strauss, J. H., & Weaver, S. C. (2001). Evolutionary relationships and systematics of the alphaviruses. Journal of virology, 75(21), 10118–10131.

20. McNamara, T. D., O’Shea-Wheller, T. A., DeLisi, N., Dugas, E., Caillouet, K. A., Vaeth, R., et al. (2021). An efficient alternative to the CDC gravid trap for southern house mosquito (Diptera: Culicidae) surveillance. Journal of Medical Entomology, 58(3), 1322–1330.

21. Cilek, J. E., Knapp, J. A., & Richardson, A. G. (2017). Comparative Efficiency of Biogents Gravid Aedes Trap, Cdc Autocidal Gravid Ovitrap, and CDC Gravid Trap in Northeastern Florida1. Journal of the American Mosquito Control Association, 33(2), 103–107.

23. Reiter, P. (1983). A portable battery-powered trap for collecting gravid Culex mosquitoes.

23. Darsie, R. F., Jr., & Ward, R. A. (2005). Identification and geographical distribution of the mosquitoes of North America, north of Mexico. University Press of Florida.

25. Thermo Fisher Scientific. (2016). TRIzol™ Reagent user guide. Thermo Fisher Scientific.

25. Unal, I. (2017). Defining an optimal cut-point value in ROC analysis: an alternative approach. Computational and mathematical methods in medicine, 2017(1), 3762651.

26. Sam, S. S., Teoh, B. T., Chee, C. M., Mohamed-Romai-Noor, N. A., Abd-Jamil, J., Loong, S. K., et al. (2018). A quantitative reverse transcription-polymerase chain reaction for detection of Getah virus. Scientific reports, 8(1), 17632.

27. Bartholomay, L. C., Waterhouse, R. M., Mayhew, G. F., Campbell, C. L., Michel, K., Zou, Z., et al. (2010). Pathogenomics of Culex quinquefasciatus and meta-analysis of infection responses to diverse pathogens. Science, 330(6000), 88–90.

29. Leta, S., Beyene, T. J., De Clercq, E. M., Amenu, K., Kraemer, M. U., & Revie, C. W. (2018). Global risk mapping for major diseases transmitted by Aedes aegypti and Aedes albopictus. International journal of infectious diseases, 67, 25–35.

29. Liu, N., Xu, Q., Li, T., He, L., & Zhang, L. (2009). Permethrin resistance and target site insensitivity in the mosquito Culex quinquefasciatus in Alabama. Journal of Medical Entomology, 46(6), 1424–1429.

30. Crockett, R. K., Burkhalter, K., Mead, D., Kelly, R., Brown, J., Varnado, W., et al. (2012). Culex flavivirus and West Nile virus in Culex quinquefasciatus populations in the southeastern United States. Journal of Medical Entomology, 49(1), 165–174.

31. Porterfield, J. S. (1975). The basis of arbovirus classification. Medical biology, 53(5), 400–405.

32. Kawamura, H., Yago, K., Narita, M., Imada, T., Nishimori, T., Haritani, M. (1987). A fatal case in newborn piglets with Getah virus infection: Pathogenicity of the isolate. The Japanese Journal of Veterinary Science, 49(6), 1003–1007.

33. Wang, F. I., Chang, C. Y., & Huang, C. C. (2019). Togaviruses. Diseases of Swine, 740–742.

34. Xing, C., Jiang, J., Lu, Z., Mi, S., He, B., Tu, C., et al. (2020). Isolation and characterization of Getah virus from pigs in Guangdong province of China. Transboundary and emerging diseases, 67(5), 2249–2253.

35. Ren, T., Mo, Q., Wang, Y., Wang, H., Nong, Z., Wang, J., et al. (2020). Emergence and phylogenetic analysis of a Getah virus isolated in southern China. Frontiers in Veterinary Science, 7, 552517.

36. ArboCat Virus_Getah (GETV). CDC, Mar. 15, 2022, https://wwwn.cdc.gov/arbocat/VirusDetails.aspx?ID=161. Accessed 15 Mar. 2024.

37. MacLachlan, N., and E. Dubovi. “Togaviridae.” Fenner’s Veterinary Virology, 4th ed., Elsevier Science & Technology, 2011, Chap. 29.

38. Brown, R. S., Wan, J. J., & Kielian, M. (2018). The alphavirus exit pathway: what we know and what we wish we knew. Viruses, 10(2), 89.

39. Lu, G., Chen, R., Shao, R., Dong, N., Liu, W., & Li, S. (2020). Getah virus: an increasing threat in China. Journal of Infection, 80(3), 350–371.

40. Li, Y. Y., Liu, H., Fu, S. H., Li, X. L., Guo, X. F., Li, M. H., et al. (2017). From discovery to spread: the evolution and phylogeny of Getah virus. Infection, Genetics and Evolution, 55, 48–55.

41. Ortiz, D. I., Piche-Ovares, M., Romero-Vega, L. M., Wagman, J., & Troyo, A. (2021). The impact of deforestation, urbanization, and changing land use patterns on the ecology of mosquito and tick-borne diseases in Central America. Insects, 13(1), 20.

42. Fournet, F., Jourdain, F., Bonnet, E., Degroote, S., & Ridde, V. (2018). Effective surveillance systems for vector-borne diseases in urban settings and translation of the data into action: a scoping review. Infectious diseases of poverty, 7(1), 99.

43. Morse, S. S. (1995). Factors in the emergence of infectious diseases. Emerging infectious diseases, 1(1), 7.

44. Gottdenker, N. L., Streicker, D. G., Faust, C. L., & Carroll, C. R. (2014). Anthropogenic land use change and infectious diseases: a review of the evidence. EcoHealth, 11(4), 619–632.

